# Impact of magnetite nanowires orientation on morphology and activity of *in vitro* hippocampal neural networks

**DOI:** 10.1101/2023.03.29.534753

**Authors:** Belén Cortés-Llanos, Rossana Rauti, Ángel Ayuso-Sacido, Lucas Pérez, Laura Ballerini

## Abstract

Nanomaterials design, synthesis and characterization are ever-expanding approaches towards developing biodevices or neural interfaces to treat neurological diseases. The ability of nanomaterials features, to tune neuronal networks morphology or functionality is still under study. In this work, we unveil how, when interfacing mammalian brain cultured neurons, iron oxide nanowires (NWs) orientation affects neuronal and glial densities, and network activity. Iron oxide NWs were synthesized by electrodeposition, fixing the diameter to 100 nm and the length to 1 μm. Scanning electron microscopy, Raman and contact angle measurements were performed to characterize the NWs morphology, chemical composition and hydrophilicity. Hippocampal cultures were seeded on NWs devices and after 14 days the cell morphology was studied by immunocytochemistry and confocal microscopy. Live calcium imaging was performed to study neuronal activity. Using random (R-NWs) a higher neuronal and glial cell densities were obtained compared with the control and vertical (V-NWs), while using V-NWs more stellate glial cells were found. R-NWs produced a reduction in neuronal activity while V-NWs increased the neuronal network activity, possibly due to higher neuronal maturity and a lower number of GABAergic neurons, respectively. These results highlight the potential of NWs manipulations to design ad hoc regenerative interfaces.

## 1. Introduction

The application of nanotechnology to nervous system repair strategies has increasingly targeted the development of nanomaterials to interface neurons and neuronal networks [1–3]. Importantly, diverse nanomaterial dimensionality can elicit distinct responses in brain cells and circuits [4,5]. In this framework, particular attention must be given to nanowires (NWs) as a permissive substrate for neural growth [6,7]. NWs nano-size enables their crossing of the plasma membrane without the emergence of significant cell damage [8]. NWs have been used for positioning neurons [9], directing neurite growth [7,10,11], and for extracellular and intracellular recording through neural cell membranes [12–15].

In the last years, NWs have been exploited in arrays by tuning their size and orientation to improve cell adhesion. Recent advances in nanotechnology have allowed further manufacturing of nanostructures with high aspect ratios, such as NWs or polymer patterns [16]. Using different fabrication technologies, novel standard integrated circuits based on NWs, namely scaffolds or multielectrode arrays (MEA), were fabricated to record the neuronal network activity [14,17]. Among these techniques, electrodeposition is an electrochemistry approach that provides precise control of NWs diameter and length by the synthesis parameters [18]. The ease of tuning NWs orientation might be relevant in the design of devices for sensing and manipulating cells [11,19,20].

Together with NWs orientation, the building material may also affect functional cellular properties, such as adhesion and survival [21–23]. High-density Silicon (Si) and Gallium Phosphide (GaP) semiconductor NWs were shown to affect the morphology of retinal or peripheral neurons [21,24,25]. In the last decades, due to their electrical and mechanical properties, carbon-based materials have been successfully used as substrates for neuronal growth for their ability to interact with neuronal membranes and synapses [26–30]. Nonetheless, depending on the material size, semiconductors and carbon-based materials might need a coating to increase their biocompatibility [31]. In this framework, iron oxide is a well-known biocompatible material [18,32,33]. Iron oxide nanoparticles (IONPs) are being used in many experimental settings in the field of regenerative medicine, including nervous systems diseases, and for cancer treatments such as hyperthermia and Magnetic Resonance Imaging (MRI) [33–37]. Among the iron oxide materials, magnetite (Fe_3_O_4_) and maghemite (γ-Fe_2_O_3_) have the particularity of being both biocompatible and biodegradable [18,33,37].

Although several studies describe the interaction between NWs and living cells, showing the impact on cell viability and network changes [7,21,22,38,39], very few investigate the ability of the NWs orientation to impact the network morphology and activity [23,25,40,41]. Taking advantage of the elongated aspect of NWs and the iron oxide properties, we studied for the first time the ability of NWs orientation to interact with an *in vitro* hippocampal network. Electrodeposited iron NWs were thermally oxidized to obtain magnetite NWs. Scanning Electron Microscopy (SEM) was used for their morphological characterization and a tensiometer for the wettability measurements. Raman Spectroscopy was performed to characterize the iron oxide present in the samples. Hippocampal cultured neurons were interfaced with the NWs, and after 8-10 Days In Vitro (DIV), we investigated cell density using immunofluorescent labelling and confocal microscopy. Live calcium imaging was used to assess the functional properties of neurons and networks. NWs orientation can affect neuronal density and glial morphology, but more interestingly, it can impact emerging synaptic network activity.

## 2. Materials and Methods

### 2.1. NWs Synthesis

Iron NWs were electrodeposited inside the nanopores of a polycarbonate membrane supplied by Sterlitech. Before the electrodeposition, an Au thin film was thermally evaporated on one side of the template to use it as a working electrode. The electrolyte was composed of 0.5 M iron sulfate heptahydrate (Fe_2_SO_4_ ·7H_2_O) and 0.5 M boric acid (H_3_BO_3_), both supplied by Panreac. The pH of the electrolyte was adjusted to a pH of 2.5 by dropping sulphuric acid (H_2_SO_4_) to the solution. The synthesis was carried out in a three-electrode vertical cell, with a Pt mesh as a counter electrode and an Ag/AgCl electrode as reference. The Fe NWs were synthesized with a constant potential of -1.15 V (vs. Ag/AgCl) being the growth rate of 12.5 nm/s. After electrodeposition, the template was removed using dichloromethane (CH_2_Cl_2_) from Sigma-Aldrich and the NWs deposited onto Si wafers or glass coverslips for characterization.

### 2.2. Physical characterization

The morphology of the samples was studied by Scanning Electron Microscopy (SEM), using a JEOL JEM 6335 microscope. The static contact angle measurements were performed using an optical tensiometer (Attension Theta, Biolin Scientific). Deionized water drops of 5 μL were deposited onto the substrate. The different angles with the baseline (left and right) and the mean value was calculated by adjusting the drop profile to a Young-Laplace curve.

Raman measurements were carried out using a Confocal Raman Microscope (Witec ALPHA 300RA) at room temperature with an Nd:YAG laser (532 nm). The spectra were measured at 0.25 mW of laser excitation power and using an objective with a numerical aperture (NA) of 0.95. Raman spectra were recorded in the spectral range of 0−3600 cm^−1^. We measured regions on the plane (XY scans, 5×5 μm^2^) and in-depth (XZ scans, 5×2 μm^2^). Punctual spectra were recorded along with the XZ, in-depth, to identify the plane with larger Raman intensity. The spectra were analyzed by using Witec Control Plus Software. The band positions were obtained fitting as Lorentzian functions.

### 2.3. Sterilization protocol

Iron NWs were deposited in glass coverslips (Kindler, EU), removing the template using dichloromethane (CH_2_Cl_2_; Sigma-Aldrich). The samples were sterilized in an oven controlling the temperature at 180 °C for 4 h to obtain magnetite NWs already sterilized. Then, 0.1 mg/mL of poly-L-lysine-coated (PLL) (Sigma-Aldrich) was dropped into the samples and left for 7 hours in the incubator at 37 °C (5% CO_2_). After that, we removed the PLL, and the coverslips were washed three times with sterilized water. The samples were left in the incubator overnight (37 °C; 5% CO_2_).

### 2.4. Preparation of primary hippocampal cultures

As previously reported, primary hippocampal cultures were prepared from postnatal day 2 or 3 (P2-P3) rats [27,42]. All procedures were approved by the local veterinary authorities and performed following the Italian law (decree 26/14) and the UE guidelines (2007/526/CE and 2010/63/UE). Animal use was approved by the Italian Ministry of Health. All efforts were made to minimize suffering and to reduce the number of animals used. All chemicals were purchased by Sigma-Aldrich unless stated otherwise. Briefly, enzymatically dissociated hippocampal neurons were plated on three different substrates: poly-L-lysine-coated, R-NWs, and V-NWs substrates at a density of 150000 cells/mL (n = 4 culture series). Cultures were incubated (37 °C; 5% CO_2_) in a medium consisting of MEM (Invitrogen), supplemented with 35 mM Glucose, 15 mM HEPES, 1 mM Apo-Transferrin, 48 μM Insulin, 3 μM Biotin, 1 mM Vitamin B12 and 500 nM Gentamicin (Gibco). The culture medium was renewed after 2 days from seeding and hereafter changed every 2 days. Cultures were then used for experiments after 8-10 DIV.

### 2.5. Immunocytochemistry and image processing

After 8-10 DIV, hippocampal cultures were washed three times with PBS and post-fixed in 4% paraformaldehyde (PFA, prepared from fresh paraformaldehyde) in PBS for 20 minutes at room temperature (RT). After that, cells were washed three times with PBS and permeabilized with 1% Triton X-100 for 30 min, blocked with 5% FBS in PBS for 30 min at room temperature, and incubated with primary antibodies for 45 min. The primary antibodies used were rabbit polyclonal anti-β-tub III (1:500 dilution) and mouse monoclonal anti-GFAP (1:500 dilution). After PBS washes, cells were incubated for 45 minutes with AlexaFluor 594 goat anti-rabbit (Invitrogen, dilution 1:500) and AlexaFluor 488 goat anti-mouse (Invitrogen, dilution 1:500). Samples were mounted in Vectashield with DAPI to stain the nuclei (Vector Laboratories) on 1 mm tick coverslips. Cells were imaged at 20 × (0.5 NA) magnification using a Leica DM6000 fluorescent microscope (Leica Microsystems GmbH, Wetzlar, Germany). Cell densities, β-tub III area, and GFAP perimeter and shape were quantified with a random sampling of seven to ten fields (713 × 533 μm^2^; control and NWs substrates, at least n=3 culture series).

For GABA staining, cells were fixed with 1% Glutaraldehyde, and 4% PFA in PBS for 1 hour in darkness, at RT. After three washes with PBS, cells were blocked with 5% BSA, 3% Triton, and 1% FBS in PBS at RT and incubated with primary antibodies for 45 min. The primary antibodies were rabbit polyclonal anti-GABA (1:300 dilution) and mouse polyclonal anti-β-tub III (1:500 dilution). After PBS washes, samples were incubated for 45 min at RT with the secondary antibodies, AlexaFluor 488 goat anti-rabbit (1:500 dilution), and AlexaFluor 594 goat anti-mouse (1:500 dilution) and DAPI for the nuclei (1:500 dilution). Samples were mounted in Vectashield on 1 mm thick coverslips. Images were acquired using a Nikon C2 confocal microscope (Nikon, Japan) at 40× magnification. Z-stacks were acquired every 500 nm, from seven to ten random fields for control and NWs substrates. Offline analysis of cell density, stellate cells, and GABAergic neurons was performed using the image-processing package Fiji. Stellate cells were defined when they possesed processes longer than their perinucelar diameters [43,44]. Offline analysis of the β-tub III area and GFAP perimeter were performed using Volocity software (Volocity 3D image and quantification analysis software, PerkinElmer, USA). The images were acquired using identical exposure settings for each set of experiments. For the β-tub III area and GFAP perimeter quantifications, the intensity threshold was defined and adjusted to 0.5 SD. The area or the perimeter within each ROI with intensity above the threshold was calculated and used for statistics.

### 2.6. Calcium Imaging

Cultures were loaded with cell-permeable Ca^2+^ dye Oregon Green 488 BAPTA-1 AM (Invitrogen). Stock solution (4 mM) of the Ca^2+^ dye was prepared in DMSO, and cultures were incubated with a final concentration of 4 μM for 30 min (37 °C; 5% CO_2_). The samples were then placed in a recording chamber mounted on an inverted microscope (Nikon *Eclipse Ti-U*) where they were continuously superfused at RT by a recording solution of the following composition (mM): 150 NaCl, 4 KCl, 1 MgCl_2_, 2 CaCl_2_, 10 HEPES, 10 glucose (pH adjusted to 7.4 with NaOH). Cultures were observed with a 20× objective (0.45 NA), and recordings were performed from visual fields (512 × 512 μm^2^, binning 4). Ca^2+^-dye was excited at 488 nm with a mercury lamp; excitation light was separated from the light emitted from the sample using a 395 nm dichroic mirror and ND filter (1/32). Images were continuously acquired (exposure time 150 ms) using an ORCA-Flash4.0 V2 sCMOS camera (Hamamatsu). The imaging system was controlled by integrating imaging software (HCImage Live). 10 μM bicuculline methiodide was bath-applied after 10 min of recording. At the end of each experiment, Tetrodotoxin (TTX, 1μM, a voltage-gated, fast Na^+^ channel blocker; Latoxan) was applied to confirm the neuronal nature of the recorded signals. Recorded images were analyzed offline with Fiji (selecting region of interest, ROI, around cell bodies) and Clampfit software (pClamp suite, 10.2 version; Molecular Devices LLC, US). Intracellular Ca^2+^ transients were expressed as fractional amplitude increase (ΔF/F_0_, where F_0_ is the baseline fluorescence level and ΔF is the rise over baseline); we determine the onset time of neuronal activation by detecting those events in the fluorescence signal that exceed at least five times the standard deviation of the noise.

### 2.7. Synchronization analysis

The mean correlation coefficient of calcium transients was computed by calculating the cross-correlation matrix between all pairs of neurons. To do so, we used the xcorr funtion in MATLAB. The xcorr function measures the similarity between the calcium activity of a neuron (i) and the lagged trace of another neuron (j) as function of the lag. The correlation coefficient is the cross-correlation value at lag = 0 at which the traces overlap perfectly. The traces of each sample were considered independently from other samples and all traces were simultaneously analyzed.

### 2.8. Data analysis and Statistics

All data are presented as mean ± SD of the mean (*n* is the number of cells, if not otherwise indicated). Statistical significance was calculated as a function of Control/R-NWs/V-NWs, and GraphPad software was used to evaluate the differeences among the group with one-way ANOVA test. A value of p < 0.05 was accepted as indicative of a statistically significant difference, and we show as *, p<0.01 as **, and p<0.001 as ***. In box plots, the thick horizontal bar indicates the median value, while the boxed area extends from the 25^th^ to 75^th^ percentiles, with the whiskers ranging from the 5^th^ to the 95^th^ percentiles.

## 3. Results

### 3.1. Synthesis and Characterization of NWs

Two different configurations were tested to address the issue of different NWs orientations on a functional brain network. Figures 1.a and 1.b show SEM images of electrodeposited iron NWs in a random (R-NWs) and vertical (V-NWs) position on the substrate. The NWs have an average diameter of 150 nm and lengths of 1μm approximately. After the thermal treatment produced during sterilization, the NWs are oxidized to iron oxide. Figures 1.c and 1.d show an optical image of R-NWs and V-NWs respectively after sterilization, where we observed the homogeneous distribution of the NWs (black) along the substrate (yellow). Wettability measurements give us the degree of wetting when a solid (NWs substrates) and liquid interact. Contact angle values for R-NWs and V-NWs were 103° and 135°, respectively. Iron oxide R-NWs present higher wettability compared with V-NWs.

**Figure 1:**
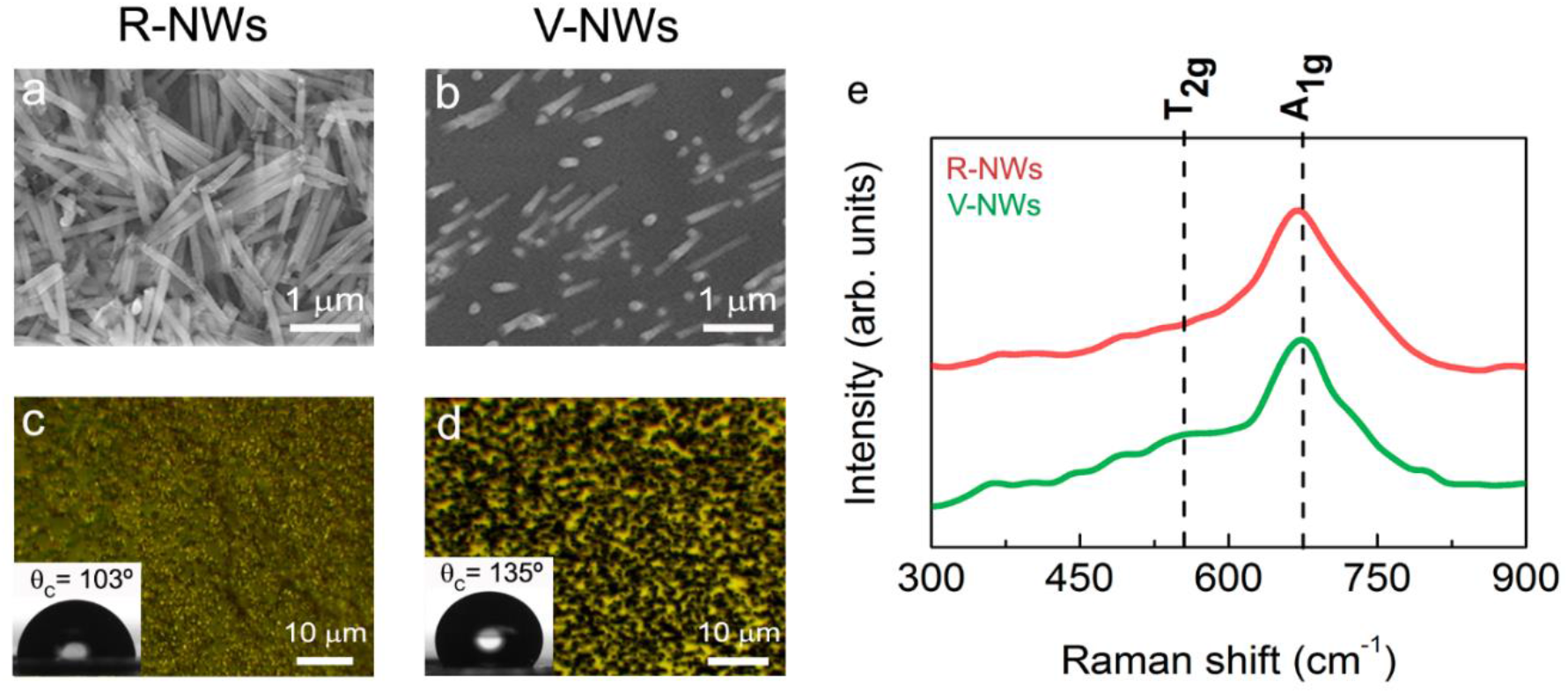
SEM images of electrodeposited iron **(a)** random NWs (R-NWs) and **(b)** vertical NWs (V-NWs). Optical images and contact angles of **(c)** R-NWs and **(d)** V-NWs orientations. **(e)** Raman spectra from 300 cm^-1^ to 900 cm^-1^ for R-NWs and V-NWs substrates. The positions for the active vibrational modes of magnetite were marked with vertical black lines. In-plane Raman intensity images measuring different single Raman spectra were taken each 100 nm with a laser excitation power of 0.25 mW for R-NWs (red) and V-NWs substrates (green).

### 3.2. Investigating hippocampal network

We compared primary hippocampal neurons upon 8-10 DIV of growth on poly-L-lysine glass coverslips (named control) with those developed on R-NWs or V-NWs substrates. Immunofluorescence techniques were used to determine the cellular composition of the different substrates by imaging the specific cytoskeletal component β-tubulin III (β-tub III) to visualize neurons (Figure 2.a-c), and glial fibrillary acidic protein (GFAP) to visualize astrocytes (Figure 2.f-h) [27,42].

**Figure 2:**
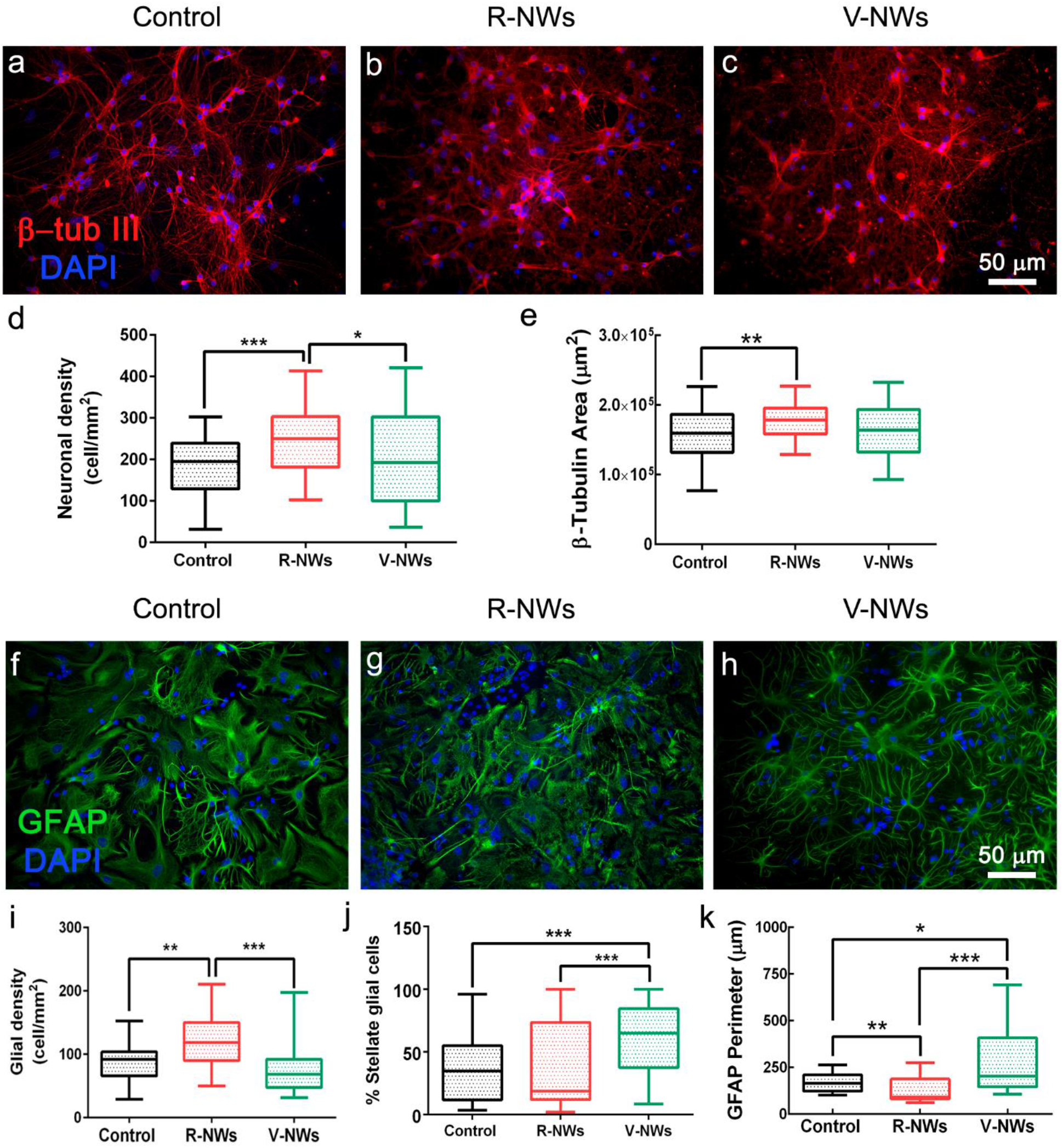
Hippocampal cell densities are regulated by NWs orientation. Immunofluorescence micrographs visualizing neurons **(a-c)**, in red anti-β-tub III, and glial cells **(f-h)**, in green anti-GFAP, in the three different conditions; nuclei are visualized by DAPI (in blue). Scale bars: 50 μm. Box plot **(d)** summarizes neuronal densities and **(e)** β-tub III area. Box plot in **(i)** summarizes glial densities, in **(j)** the percentage of stellate glial cells, and in **(k)** the GFAP perimeter for the substrates tested.

We studied the neuronal density (estimated by quantifying β-tub III cells) among the three different growth conditions. We measured 182.4 ± 73.9 neurons/mm^2^ in control (n = 59 visual fields, 8 samples, n = 3 series of cultures), in R-NW 241.2 ± 74.4 neurons/mm^2^ (n = 62 visual fields, 7 samples, n = 3 series of cultures) and V-NWs 202.6 ± 115.0 neurons/mm^2^ (n=59 visual fields, 8 samples, n= 3 series of cultures) (Figure 2.d), with a significant increase in cell density in R-NWs substrates (***p < 0.001 and *p < 0.05, Kruskal-Wallis, one-way ANOVA) when compared to control and to V-NWs cultures. Consistently, we observed a significant increase (**p < 0.01, Kruskal-Wallis, one-way ANOVA) in β-tub III area for R-NWs ((1.75 ± 0.24) · 10^5^ μm^2^, n = 69 visual fields, 7 samples) when compared with control ((1.58 ± 0.36) · 10^5^ μm^2^, n = 65 visual fields, 8 samples). V-NWs displayed lower values of β-tub III area ((1.58 ± 0.37) · 10^5^ μm^2^, n = 68 visual fields, 8 samples) when compared to R-NWs, although without reaching statistical significance (Figure 2.e).

With a similar approach, we investigated the effect of R-, V-NWs substrates on glial cell density. We identified and quantified GFAP^+^ cells in control (87.4 ± 26.8 GFAP cells/mm^2^, n = 45 visual field, 6 samples), in R-NWs (119.1 ± 40.4 GFAP-cells/mm^2^, n = 46 visual field, 5 samples) and V-NWs (73.7 ± 34.8 GFAP-cells/mm^2^, 4 visual field, 6 samples). We observed a significant increase of GFAP^+^ cells in R-NWs substrates when compared with control (**p<0.01, Kruskal-Wallis, one-way ANOVA) and with V-NWs (***p< 0.001, Kruskal-Wallis, one-way ANOVA) (Figure 2.i). We further quantified the presence of glial cells displaying a stellate morphology (see methods) as a percentage of stellate glial cells on the total amount of GFAP^+^ cells. We detected (36.7 ± 25.9)% of stellate GFAP^+^ cells in the control (n = 43 visual fields, 6 samples), similarly in R-NWs (38.2 ± 32.3)%, (n = 44 visual fields, 5 samples), while stellate morphology ratio was significantly increased by V-NWs substrates (61.7 ± 26.0)%, n = 43 visual fields, 6 samples; (***p < 0.001, Kruskal-Wallis, one-way ANOVA) (Figure 2.j). To further assess GFAP^+^ cells features, we calculated the GFAP perimeter (Figure 2.k). The values detected in control (161.3 ± 48.1 μm, n = 51 visual fields, 6 samples), R-NWs (117.6 ± 59.6 μm, n = 50, 5 samples), and V-NWs (237.4 ± 165.8 μm, n = 44, 6 samples) conditions are in accordance with the ratio of stellate glial cells detected with high values in V-NWs (*p < 0.05, Kruskal-Wallis, one-way ANOVA vs. control and ***p < 0.001, Kruskal-Wallis, one-way ANOVA vs. R-NWs).

### 3.3. Exploring neuronal activity

To investigate the network dynamics of neuronal cells grown on R- and V-NWs, we monitored neuronal emerging activity using fluorescence calcium imaging (see methods) of representative fields (318.2 × 318.2 μm^2^). At 8-10 DIV, neurons are synaptically connected and display spontaneous activity, including irregular bursts of synchronized firing epochs [27,28,42]. Figures 3.a-c snapshots show the recorded fields and spatial distribution of recorded cells simultaneously traced within the same field of view, with single-cell resolution. We compared and characterized spontaneous Ca^2+^ oscillations (Figure 3.d); such activity was always fully blocked by tetrodotoxin (TTX, 1 μM) application [27,46].

**Figure 3:**
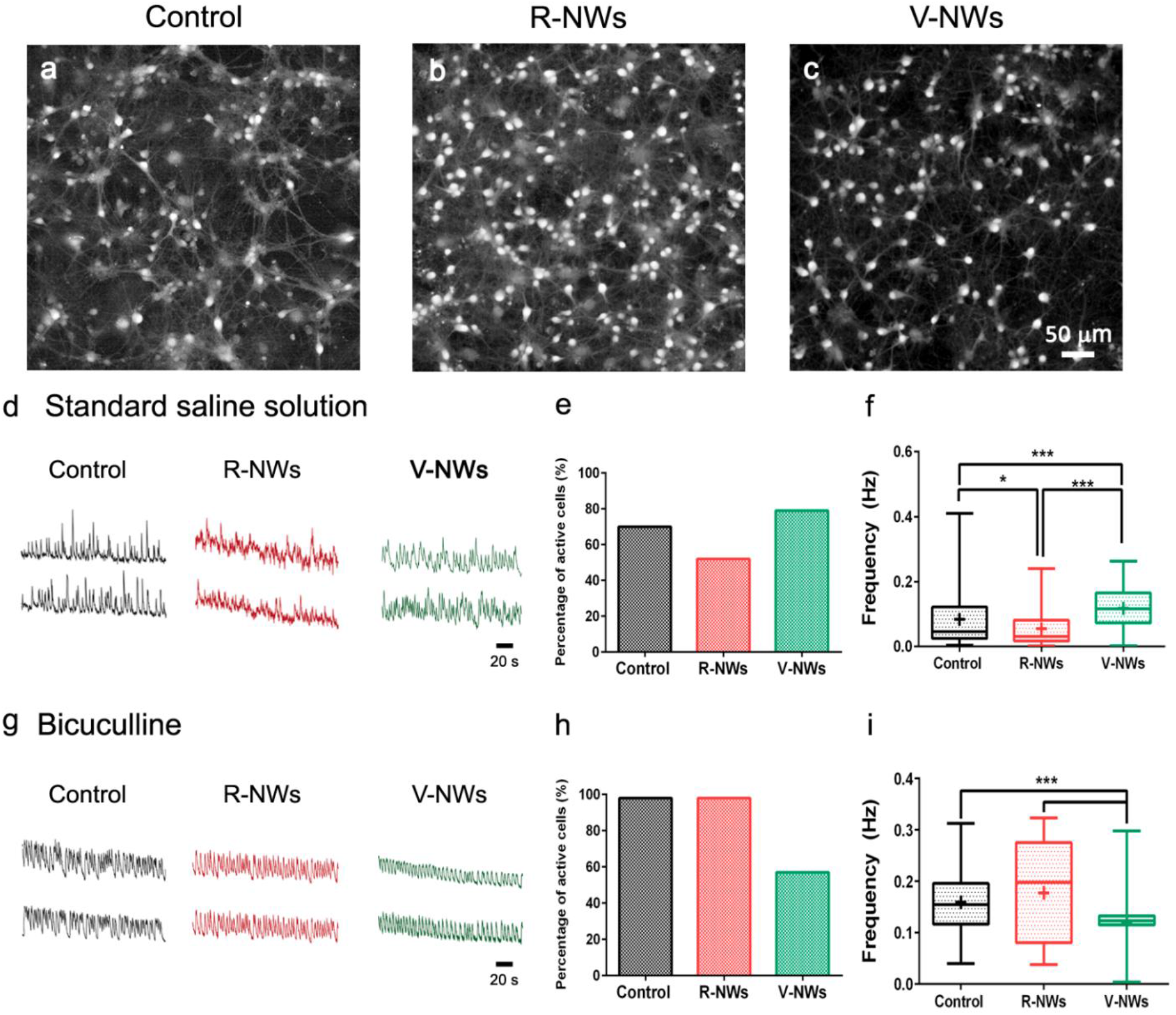
Hippocampal network activity is regulated by NWs orientation. Snapshots of representative fields of neuronal cultures grown on control in **(a)**, R-NWs in **(b)**, and V-NWs in **(c)** substrates cells are visualized by Oregon-Green 488 BAPTA-1 AM. Representative fluorescence tracings of repetitive Ca^2+^-transients in standard saline solution **(d)** or bicuculline-induced **(g)** recorded in hippocampal cultures of 8-10 DIV grown on control (black), R-NWs (red) and V-NWs (green) (two-sample neurons were selected from the same field). Bar plots summarize the percentage of spontaneously active cells (in **e** and **h**, standard saline solution and bicuculline, respectively), and box plots frequencies values for (**f** and **i**; in standard saline solution and bicuculline, respectively).

In our recordings, spontaneous Ca^2+^ activity was detected in 70% of cells grown on glass coverslips, 51% on R-NWs and 79% on V-RNWs, detecting no significant differences between control and magnetite substrates (Fisher’s exact test, n = 8 visual fields for control, n = 6 R-NWs and n = 6 V-NWs, n = 3 series of cultures) (Figure 3.e). We measured the occurrence of spontaneous Ca^2+^ episodes in active cells by quantifying the frequency distribution that was significantly (*p < 0.05, Kruskal-Wallis, one-way ANOVA) reduced in R-NWs (0.034 ± 0.051 Hz, n = 95 cells, 5 different samples) when compared to control ones (0.051 ± 0.088 Hz, n = 273 cells, 8 different samples; plot in Figure 3.f). Conversely, in V-NWs, the frequency of calcium events was significantly higher (0.096 ± 0.058 Hz, n = 564 cells, 6 different samples; ***p < 0.001, Kruskal-Wallis, one-way ANOVA) when compared to both controls and R-NWs (Figure 3.f), despite the lower neuronal density compared to R-NWs).

In the second set of experiments, we pharmacologically block GABA_A_ receptors by bicuculline (10 μM; 20 min) application. In Figure 3.g, fluorescent tracings show the appearance of Ca^2+^ episodes brought about by bicuculline in active cells. Upon this treatment, the percentage of active cells increased in all conditions without significant differences between the three substrates: (98 ± 2)% in control (n = 7), (98 ± 2)% in R-NWs cultures (n = 4), and (57 ± 38)% (n = 5) in V-NWs ones (Figure 3.h). Moreover, the frequency of calcium events increased to 0.148 ± 0.058 Hz (n = 430 cells from 7 different cultures) in control cultures, not significantly different when compared to R-NWs cultures (0.148 ± 0.097 Hz; n = 119 cells from 5 different cultures) but was significantly (***p < 0.001, Kruskal-Wallis, one-way ANOVA) higher when compared to V-NWs (0.110 ± 0.030 Hz).

In summary, regardless of the diverse neuronal cell density, we did not detect differences in the number of active cells among the three substrates. Nevertheless, cells in V-NWs were significantly more active when compared to control and R-NWs in standard medium. Still, the difference was reverted upon pharmacological removal of GABA_A_ receptor-mediated activity. This prompts us to investigate and compare the number of GABAergic neurons among all conditions to inspect the network contribution of this cell phenotype. Neurons were co-immunostained with antibodies for β-tub III and anti-GABA (Figure 4). We identified GABAergic cells as double immunolabeled neurons. We detected a significant (***p < 0.001, Kruskal-Wallis, one-way ANOVA) reduction in their number in V-NWs cultures ((9 ± 5)%, n = 29 visual fields from 3 different cultures) compared to control ((15 ± 9)%, n = 30 visual fields from 3 different cultures; see the plot in Figure 4.d). No significant differences were found comparing control or V-NWs cultures with R-NWs ones ((12 ± 9)%, n = 28 visual fields from 3 different cultures).

**Figure 4:**
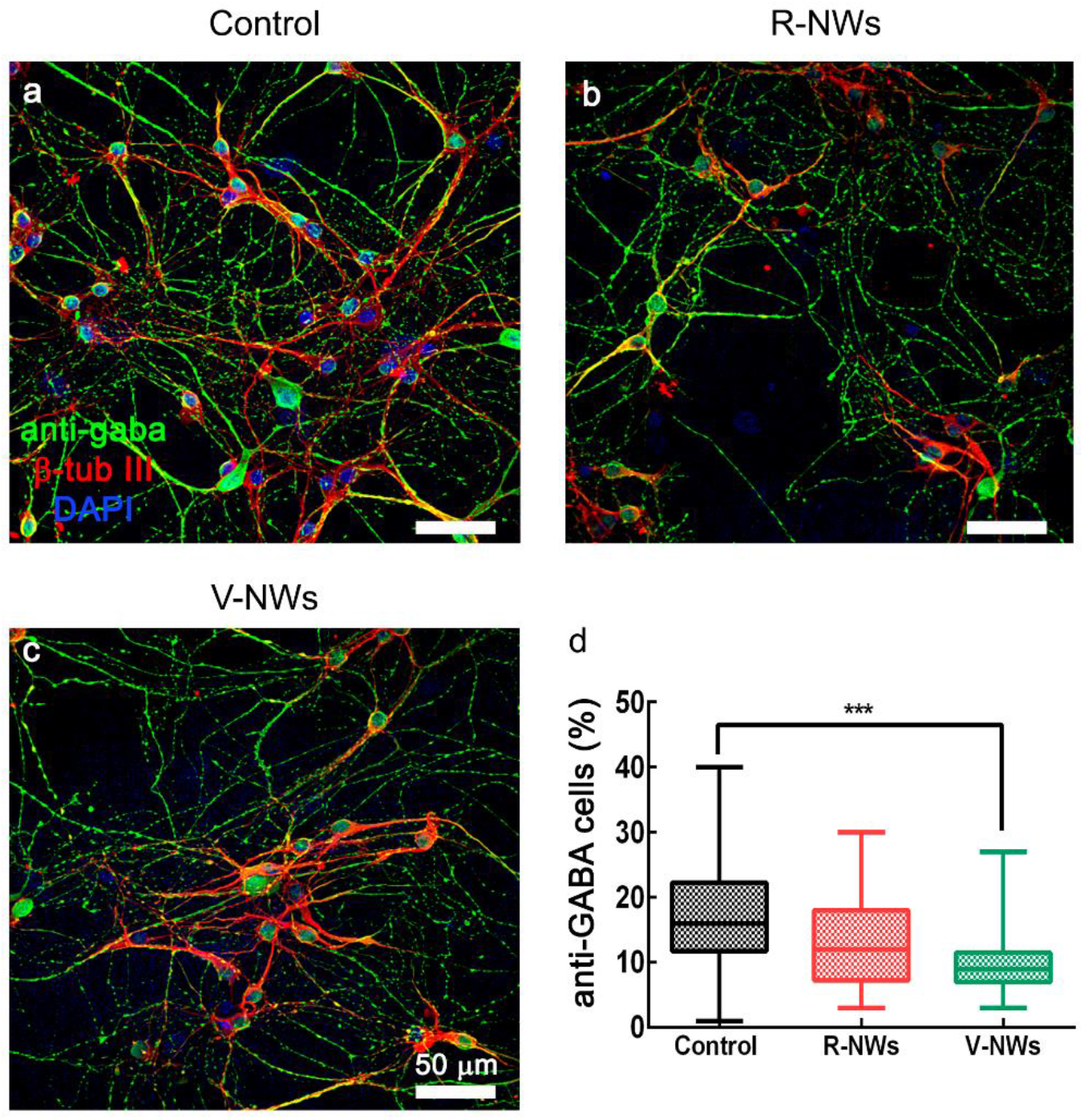
GABAergic neuronal phenotype is regulated by NWs orientation. Confocal micrographs show hippocampal cultures grown (8-10 DIV) on **(a)** Control, (b) R-NWs and **(c)** V-NWs magnetite platforms immune-stained for GABA (in green), β-tub III (in red) and DAPI (in blue). Scale bar: 50 μm. (d) The plot summarizes the percentage of GABA-ergic positive neurons.

## 4. Discussion

We report here the ability of NWs orientation to impact with neuronal growth and functionality. Nanowires’ biocompatibility depends on their diameter, length (a few microns), and the building material [21,22,47]. Furthermore, studies proved Fe_3_O_4_ biocompatibility [32]. Fe_3_O_4_ NWs platforms, with lengths of 1 μm will not present any viability issue. The differences regarding the contact angle values between R-NWs and V-NWs were due to their orientation. Orientated NWs substrates (V-NWs) possess contact angle values closer to superhydrophobic materials, which contact angle values of about 150°, due to their higher aspect ratio [48]. These changes in the aspect ratio could affect cell adhesion and its network.

We observed that NWs orientation plays an important role in modulating the morphology of the neural cellular network. Accordingly, when we used V-NWs, the morphology of the neural cell network was not affected, confirming that the orientation of NWs impacts cell adhesion. In the same line, the neuronal area (β-tub III area) was increased in R-NWs substrates, indicative of higher neuronal branching [49]. This effect might be related to adhesion processes [49]. Studies showed that using InAs NWs or Si NWs as a substrate, it was possible to obtain higher neurite outgrowth or longer neurites than in a planar control [23,50]. However, we did not observe differences in β-tub III cells between V-NWs and the control substrates due to the high hydrophobic surfaces presented on V-NWs compared with R-NWs. Therefore, using NWs substrates can increase the β-tub III area, and this parameter was associated with the orientation. We obtained a higher β-tub III area using R-NWs than on V-NWs compared to the control. In accordance with the higher cell adhesion promoted by R-NWs is the increased number of glial cells (GFAP^+^). Intriguingly, the morphology of GFAP^+^ cells was also affected by NWs; in the V-NWs platform, besides reducing glial cell adhesion compared with R-NWs, an interesting feature when developing electrodes for brain implants; [51] augmented the number of stellate shape astrocytes. Substrates with higher thickness can favour this stellate shape on glial cells since they can simulate an *in vivo* model [5,52]. Previous investigations showed that the cell volume and processes morphology can change astrocyte function [53]. Furthermore, astrocyte morphology is key to their function and communication with neurons [54]. Astrocyte two-dimensional morphology *in vitro* becomes stellate when β-adrenergic receptors are activated [55]. It is plausible that these morphological changes are associated with the changes in network signaling we observed in V-NWs platforms.

In this study the orientation of NWs in the platform induced a different response in the neuronal activity in terms of frequency. Neuronal networks interfaced with V-NWs displayed high spontaneous activity when recorded in live imaging, compared with the control and the R-NWs. In R-NWs substrates we obtained lower frequency of neuronal activity in the standard saline solution, increased later by bicuculline-treatment, namely by removal of synaptic inhibition, reaching the control frequency values. To understand the increase in activity frequency using V-NWs and the decrease with R-NWs platform, we calculated the mean correlation coefficient related to neuronal synchronization. This analysis showed no statistically differences in the traces of syncronization between the control, R-NWs and V-NWs (Figure S1). These changes in neuronal activity might be related to different network compositions due to growth susbtrates, in particular in the amount of GABAergic neurons. These cells are in charge of the inhibition of the neuronal network, and we found out that V-NWs possess a lower amount of GABAergic neurons compared with the control, in accordance with the fact that the frequency on V-NWs substrates was not affected by bicuculline. There were no significant differences between the control and R-NWs substrates in the amount of GABAergic neurons, but in R-NWs we observed an increase in the frequency when we applied bicuculline, most probably due to a different level of maturation of synaptic inhibition on R-NWs substrates. In this regard, a previous study showed that InAs NWs substrates orientation influences cell maturation [50].

## 5. Conclusions

In summary, here we found that the orientation of NWs in the platforms is important and can modulate the morphology and activity in the interfaced neuronal network. R-NWs produced a higher area of β-tub III. On the contrary, V-NWs platforms produced the same amount of β-tub III as the control, suggesting that the NWs orientation affects cell adhesion. In terms of neuronal activity, R-NWs had a decrease in neuronal activity frequency. However, V-NWs substrates produced an increase in neuronal activity, probably due also to a change in GFAP cell morphology, confirming the ability of NWs orientation to impact neuronal network properties. NWs orientation regulated also GABAergic neuronal phenotype, with fewer GABAergic cells contributing to network excitability. Future efforts should focus on the interface interaction between the NWs and the cell membrane by high-quality live imaging and transmission electron microscopy.

## Supplementary Materials

Figure S1: Synchronization analysis.

## Author Contributions

Conceptualization, B. C-L. and L.B.; methodology, B.C-L., R.R.; formal analysis, B.C-L.; investigation, B.C-L.; resources, L.P., A.A-S., L.B.; data curation, B.C-L.; writing—original draft preparation, B.C-L., R.R.; writing—review and editing, B.C-L., R.R., L.P., A.A-S., L.B.; visualization, B.C-L.; supervision, R.R., L.B.; project administration, B.C-L.; funding acquisition, L.P., A.A-S., L.B. All authors have read and agreed to the published version of the manuscript.

## Data Availability Statement

All the data needed to evaluate the conclusions in the paper are present in the paper and/or the Supplementary Materials. Correspondence and requests for materials should be addressed to B.C.L., R.R. and L.B.

## Acknowledgments

We thank to F.P. Ulloa Severino for sharing the MATLAB code for the correlation analysis. A. del Campo from ICV-CSIC for the Raman spectra measurements.

