## Supplementary Information for "Impact of magnetite nanowires orientation on morphology and activity of *in vitro* hippocampal neural networks"

[1] Departamento de Fisica de Materiales, Universidad Complutense de Madrid, 28040, Madrid, Spain..
[2] Fundación IMDEA Nanociencia, C/ Faraday 9, 28049, Madrid, Spain.
[3] Department of Medicine, Duke University, 27705, Durham, North Carolina, USA
[4] International School for Advanced Studies, SISSA, 34136, Trieste, Italy..
[5] Deparment of Biomolecular Sciences, Università degli Studi di Urbino Carlo Bo, 61029, Urbino, Italy..
[6] Brain Tumor Laboratory, Fundación Vithas, Grupo Hospitales Vithas, 28043, Madrid, Spain..
[7] Faculty of Experimental Science and Faculty of Medicine, University of Francisco de Vitoria, 28223, Madrid, Spain.

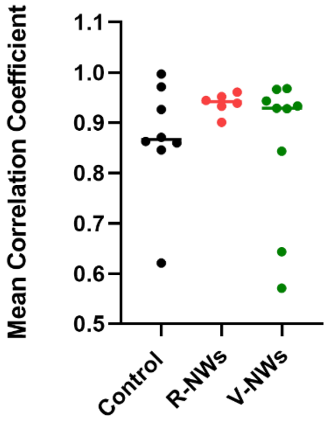


**Figure S1:** Synchronization analysis. Mean correlation coefficient of calcium transients in cultured neurons in a control, R-NWs, and V-NWs substrate.
